# Combinatorial Library Designing and Virtual Screening of Cryptolepine Derivatives against Topoisomerase II by Molecular Docking

**DOI:** 10.1101/2020.06.08.141176

**Authors:** Maria, Zahid Khan

**Affiliations:** Institute of Chemical Sciences, University of Peshawar, Pakistan

## Abstract

Computational approaches have emerging role for designing potential inhibitors against topoisomerase 2 for treatment of cancer. TOP2A plays a key role in DNA replication before cell division and thus facilitates the growth of cells. This function of TOP2A can be suppressed by targeting with potential inhibitors in cancer cells to stop the uncontrolled cell division. Among potential inhibitors cryptolepine is more selective and has the ability to intercalate into DNA, effectively block TOP2A and cease cell division in cancer cells. However, cryptolepine is non-specific and have low affinity, therefore, a combinatorial library was designed and virtually screened for identification of its derivatives with greater TOP2A binding affinities.

A combinatorial library of 31114 derivatives of cryptolepine was formed and the library was virtually screened by molecular docking to predict the molecular interactions between cryptolepine derivatives and TOP2A taking cryptolepine as standard. The overall screening and docking approach explored all the binding poses of cryptolepine for TOP2A to calculate binding energy. The compounds are given database number 8618, 907, 147, 16755, and 8186 scored lowest binding energies of −9.88kcal/mol, −9.76kcal/mol, −9.75kcal/mol, −9.73kcal/mol, and −9.72kcal/mol respectively and highest binding affinity while cryptolepine binding energy is −6.09kcal/mol. The good binding interactions of the derivatives showed that they can be used as potent TOP2A inhibitors and act as more effective anticancer agents than cryptolepine itself. The interactions of derivatives with different amino acid residues were also observed. A comprehensive understanding of the interactions of proposed derivatives with TOP2A helped for searching more novel and potent drug-like molecules for anticancer therapy. This Computational study suggests useful references to understand inhibition mechanisms that will help in the modification of TOP2A inhibitors.

## Introduction

Cancer is a life threatening, second deadliest disease causing about 1 in every 7 deaths worldwide, more than Acquired Immune Deficiency Syndrome (AIDS) [1]. The chances of cancer will increase from 14 million to 22 million in the next two decades [2]. The total medical expenditure for cancer in the US in 2013 was $74.8 billion [3].

Cancerous cells with accelerated rate undergo mitosis, transcription and replication of Deoxyribonucleic acid (DNA). Enzyme inhibition by drug is one of the potent factors to cease the uncontrolled cell growth [4]. These drugs induce conformational changes in enzymes [5]. Topoisomerase II alpha (TOP2 enzyme) maintains the integrity of genome by helping in DNA replication, transcription and chromosome segregation by unwinding the double strands of DNA to further encourage the life maintaining process of cell growth [6]. TOP2 is an effective target for cancer treatment as it induces DNA damage, by making complex with highly effective anticancer drugs. TOP2 poison blocks transcription and replication by formation of many DNA strands breaks and stop the uncontrolled division of abnormal cells which subsequently commit apoptosis [7].

Cryptolepine is an active alkaloid and has many useful properties due to its ability to intercalate into DNA strands, inhibit TOP2 and further stop DNA synthesis [8]. Cryptolepine is a very commonly used inhibitor to study apoptotic processes and its structure can be easily manipulated to make it ideal template for pro-apoptotic activity [9].

Cryptolepine has been proven to have excellent TOP2 inhibition activity, but due to its low specificity it is not used as an anticancer drug. However, there is huge potential in this compound to be converted into effective anticancer drug by making its derivatives. Addition of small functional groups to cryptolepine at various positions will result in derivatives with altered activities [10]. Synthesis and screening of these large number of derivatives is a very expensive, laborious and time consuming procedure. Computer assisted drug designing tools have proven to be successful in preliminary screening of computational library of potential candidate molecules. From the library of millions of molecules a set of few potential active molecules is predicted which can later be verified by *in vivo* and *in vitro* activity assays [11].

## Methodology

The major steps involved are:

### 3.1 Retrieval and Refinement of Crystal Structure

The crystal structure of receptor TOP2A was downloaded from protein data bank (PDB) with accession code of 5GWK. The crystal structure was refined and its missing part was modeled through Modeller software. The receptor molecule was then energy minimized using CHARMM-27force field implemented in MOE software [12].

### 3.2 Combinatorial Library Designing

Combinatorial library designing is a newly leading field that can be used to discover a group of best substituents with enhanced potencies which will help in discovering the lead compounds [13]. The combinatorial library of 31114 molecules was designed using cryptolepine as template shown in Figure 3.1 by Chem-T software [14].

**Figure 3. 1:**
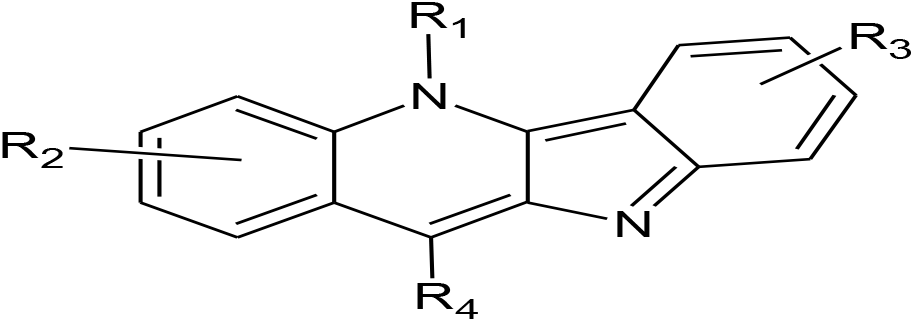
Structure of cryptolepine with R_1_, R_2_ (-CH_3_, -CH_2_CH_3_, -C_3_H_5_, -C_6_H_5_, -C_6_H_4_CH_3_, -C_6_H_4_OH, -C_6_H_4_Br) R_3_ (-CH_3_, -OCH_3_, -Cl, -F, -I) and R_4_ (-CH_3_, -C_3_H_5_, -C_6_H_4_CH_3_) indicating positions for the addition of various small substituents

For modification, four positions were selected in the template ligand structure as shown by the R-Groups in the above figure [fig 3.1]. The 2D structure of template ligand cryptolepine and the list of desired functional groups to be placed at four R-positions were drawn through Chem sketch software and were saved in mol and smile format respectively.

The template structure and the functional groups were uploaded in the Chem-T software. This was followed by the generation of 3D ligand library of 31114 molecules in the mol2/ sdf format. The molecules were energy minimized through MOE using MFF94X force field and finally visually inspected for defects [15].

### 3.3 Docking and Virtual Screening

Molecular docking is used in drug discovery process to predict the lead compounds by evaluating the protein-ligand interactions details. The molecular docking process depends on two approaches.

First, it estimates protein-ligand binding affinity on the basis of force-fields. Secondly, it searches the conformational space for protein-ligand different binding poses [16]. In docking the library is screened for the prediction of lead compounds. Molecular docking is shown graphically in Figure 3.2.

**Figure 3. 2:**
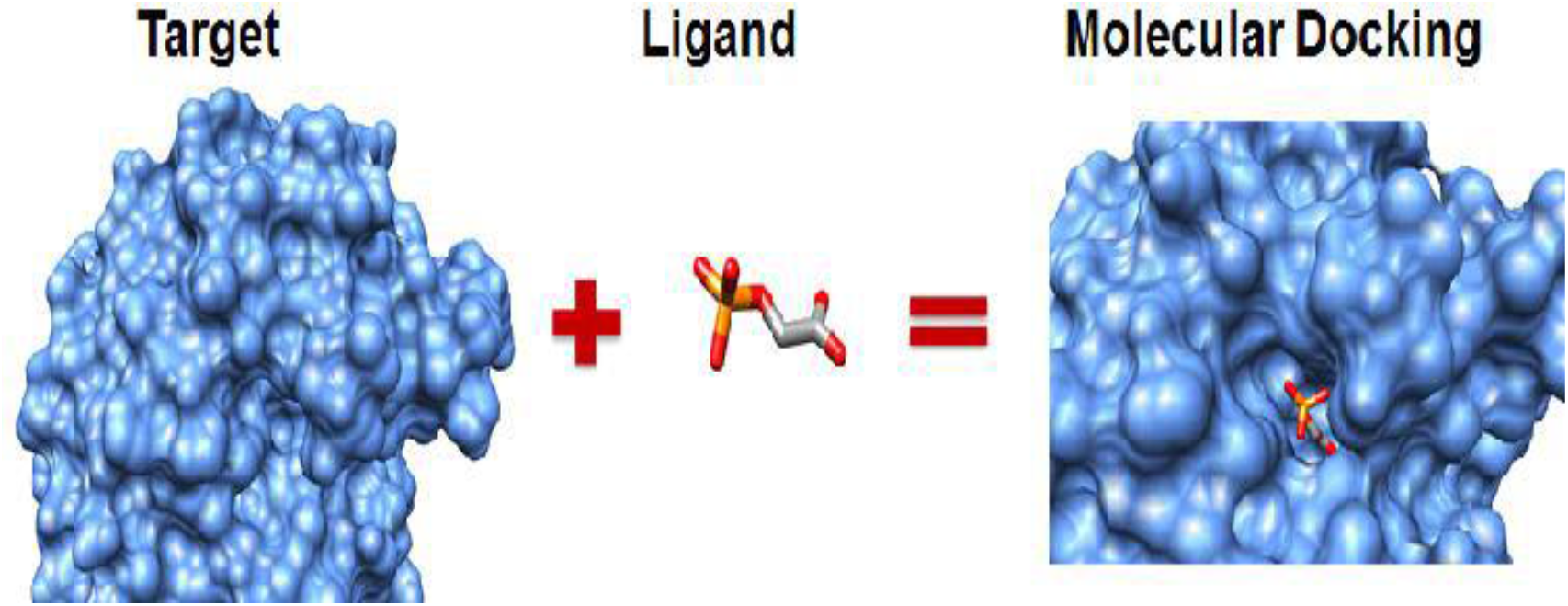
Structure showing the concept of molecular docking

Virtual screening can be defined to evaluate the library of large number of compounds for the prediction of optimized candidate compounds using different computer-based tools. It works on the basis of protein-ligand binding interactions and scoring functions. Figure 3.3 shows the binding of Ligand with receptor. Also Binding pocket of TOP2A for cryptolepine is shown in Figure 3.4 below.

**Figure 3. 3:**
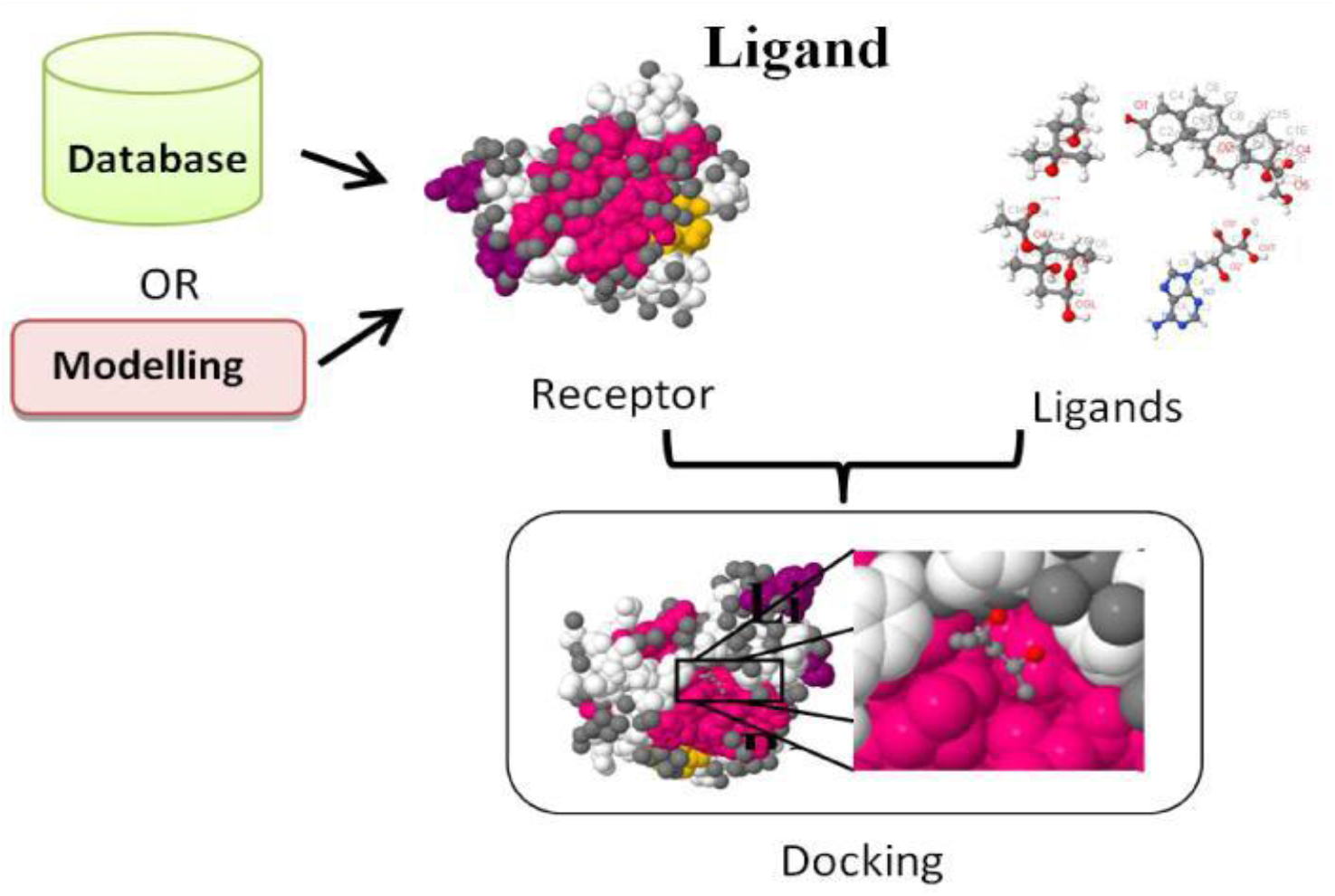
Illustration of docking of many ligands with the receptor.

**Figure 3. 4:**
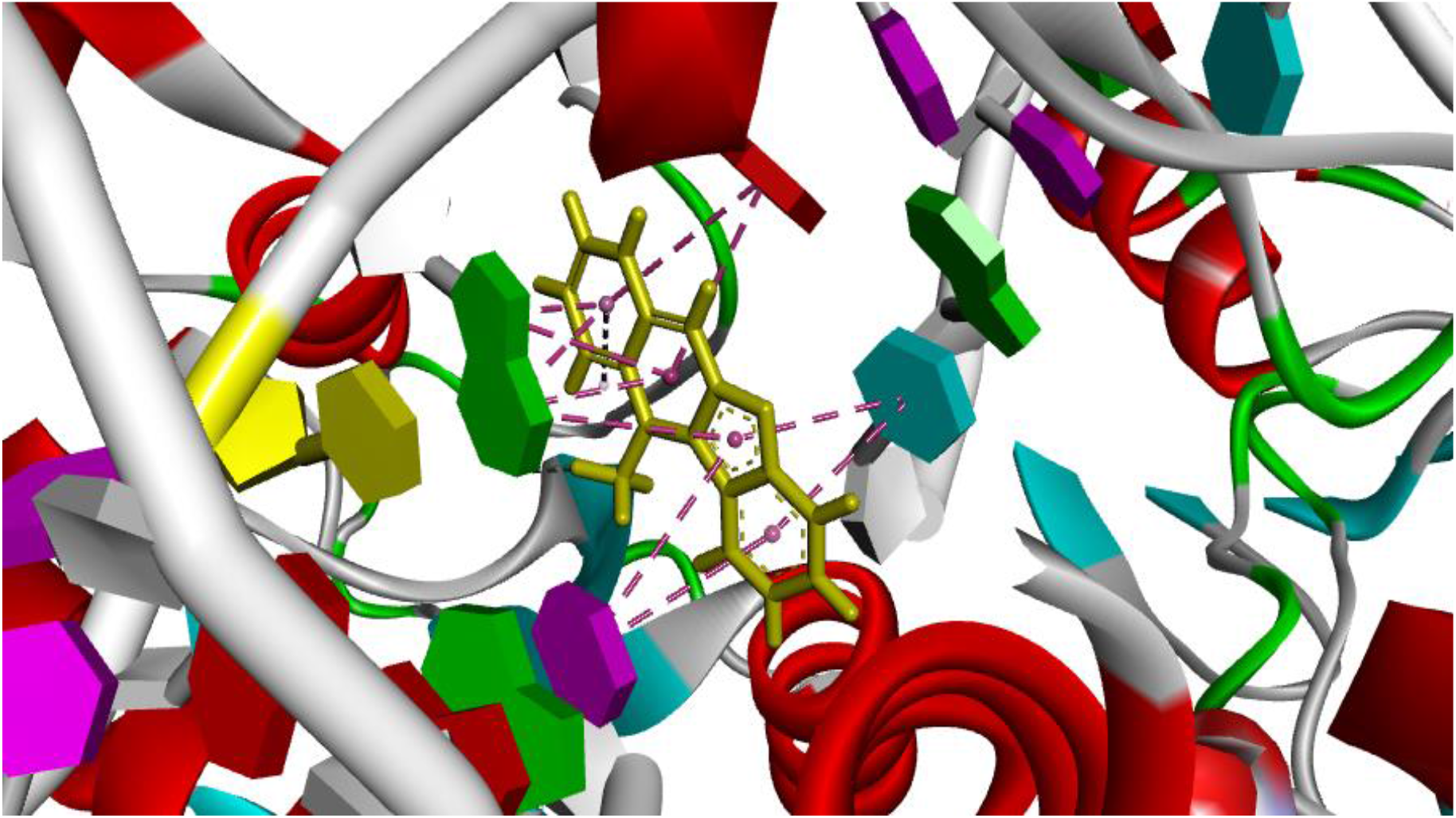
Structure of cryptolepine within binding pocket of TOP2A

### 3.4 Docking Protocol applied by MOE

Different computer based-tools are used for molecular docking. Among them, MOE is used for *in-silico docking* and virtual screening. This software is the most appropriate and reliable using accurate force-field and other docking parameters for virtual screening.

The molecules are get prepared for docking by the removal of water molecules and addition of hydrogen atoms from the 3D reported crystal structure of TOP2A (5GWK). The receptor macromolecule and library of around 31114 ligand molecules were set for docking. For reproducibility and calibration of docking software other reported inhibitors of TOP2A were also prepared in the same way. Then these ligands were also docked. Receptor macromolecule and the ligand molecules were then loaded in the MOE software for docking.

Ligand conformations were generated with the bond rotation method. These were then placed in the site with the Triangular Matcher Method, and ranked with London dG scoring function. The Retain option specifies the number of poses to pass to Refinement, for energy minimization in the pocket, before rescoring with GBVI / WSA dG scoring function [14].

### 3.5 Docking Analysis

Docking simulation analysis was carried out through discovery studio visualizer software [15]. Interaction analysis i.e., dipolar interactions, electrostatic interactions, intermolecular forces, dihedral angles, hydrogen bonding, and hydrophobicity were carried out through Visual Molecular Dynamics (VMD) and Discovery studio visualizer. Among the library of the derivatives, the best-docked compounds having a greater binding affinity for enzyme were considered as lead compounds.

## 4. Results and Discussions

### 4.1 Combinatorial Library Designing and Docking Analysis

TOP2A (5GWK) is an effective enzyme for anticancer drug target for cancer treatment. Using MOE docking software 31114 cryptolepine derivatives were virtually screened for identification of novel and potent inhibitors of TOP2A. Among 31114 compounds only five compounds were taken. And the best binding affinity and specificity of that potent top 5 derivatives were predicted and calculated by MOE software.

### 4.2 Etoposide as co-crystalized inhibitor of TOP2A

In docking studies the co-crystalized ligand etoposide in the template structure of TOP2A (5GWK) was used to show the residues involved in the active site of TOP2A as there is no co-crystalized structure of cryptolepine with TOP2A. The amino acid residues present in the binding pocket of TOP2A include Met 1083, Gly 805, Gly 779, Arg 804, Asp 780 and Met 1079, while nucleotides involved are dA 1516, dG1517, dG 1534, dC 1532 and dT 1533.

**Figure 4. 1:**
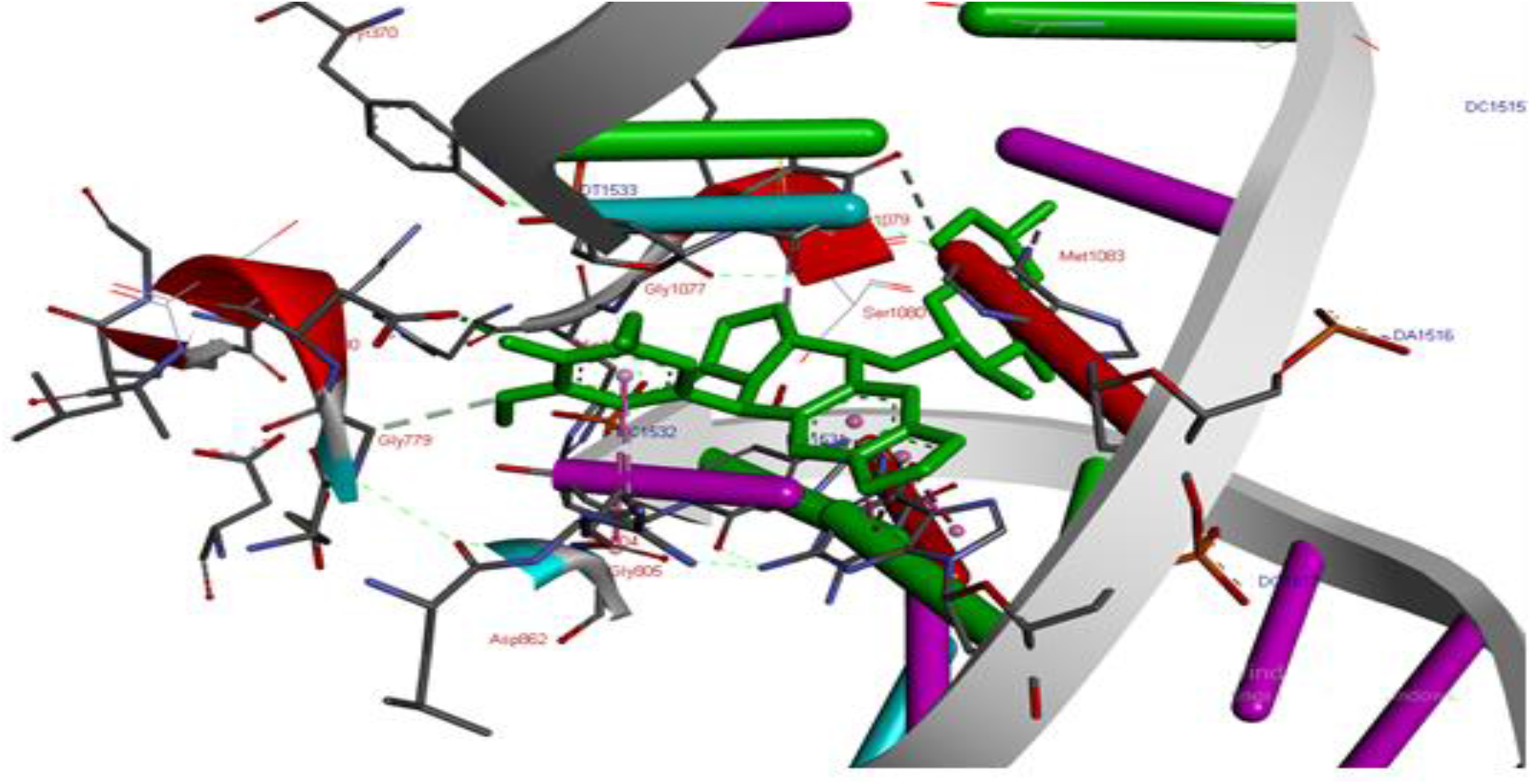
3D structure showing etoposide and its interacting residues in binding pocket of TOP2A through Discovery Studio Visualizer

**Figure 4. 2:**
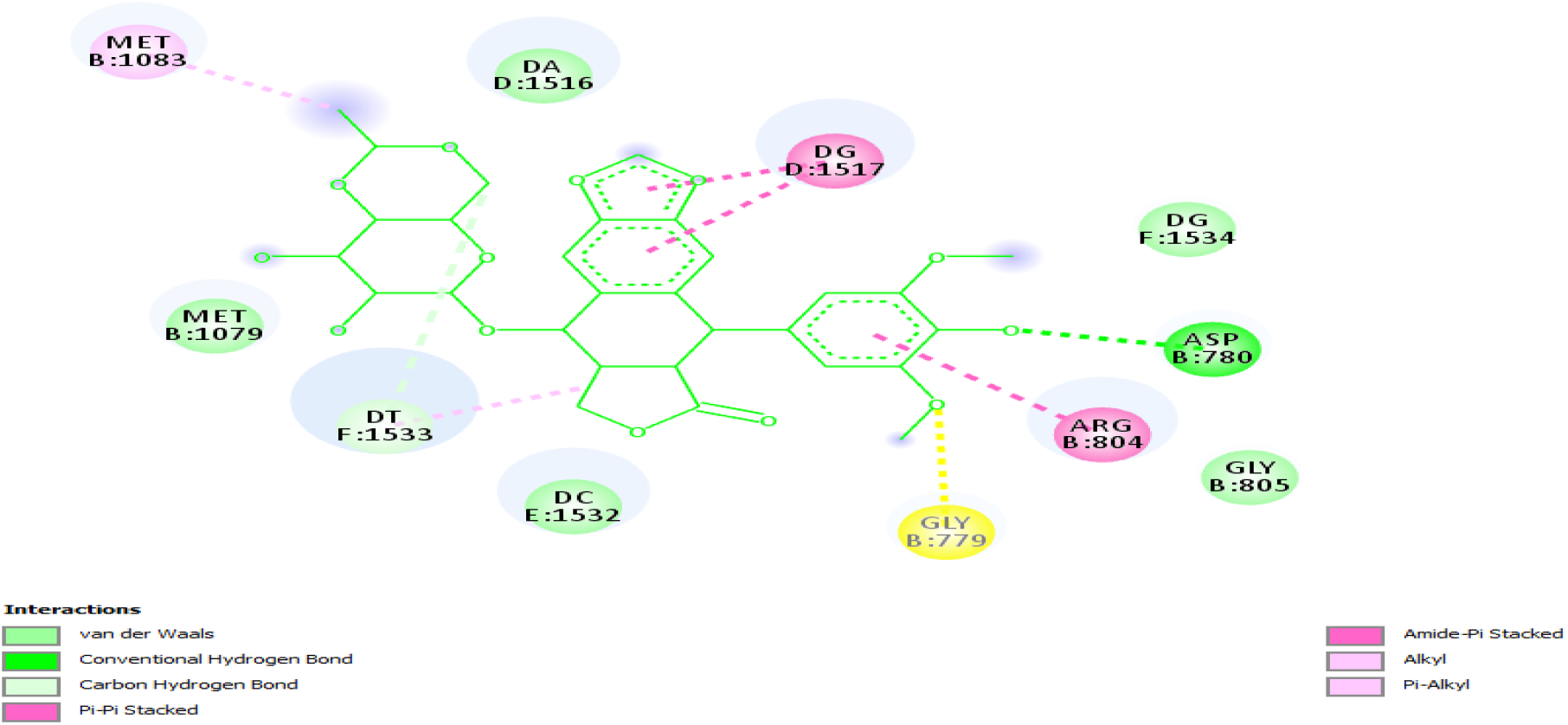
2D structure of Etoposide representing the interacting residues with the receptor

The above structure of etoposide shows the interacting residues with the receptor and form non hydrophbic interactions such as hydrogen bond, van der waals, carbon hydrogen bond, pi-pi stacked, amide-pi stacked, alkyl and pi-alkyl.

### 4.3 Cryptolepine as parent inhibitor for TOP2A

Cryptolepine was used as parent compound to target the unwinding property of TOP2A during replication and transcription of DNA. The amino acid residues in the binding pocket of co-crystalized TOP2A include Met 1079, Gly 805, Arg 804 and the double stranded DNA purine- pyrimidine such as dC 1532, dT 1533, dA 1516 and dG 1517.

Different functional groups attached with cryptolepine at **R_1_**, **R_2_** are -CH_3_, -CH_2_CH_3_, -C_3_H_5_, -C_6_H_5_, - C_6_H_4_CH_3_, -C_6_H_4_OH, -C_6_H_4_Br, **R_3_** has -CH_3_, -OCH_3_, -Cl, -F, -I and **R_4_** has -CH_3_, -C_3_H_5_, -C_6_H_4_CH_3_. The positions of **R_1_, R_2_, R_3_** and **R_4_** are distributed over the rings as shown in the below Figure 4.3.

**Figure 4. 3:**
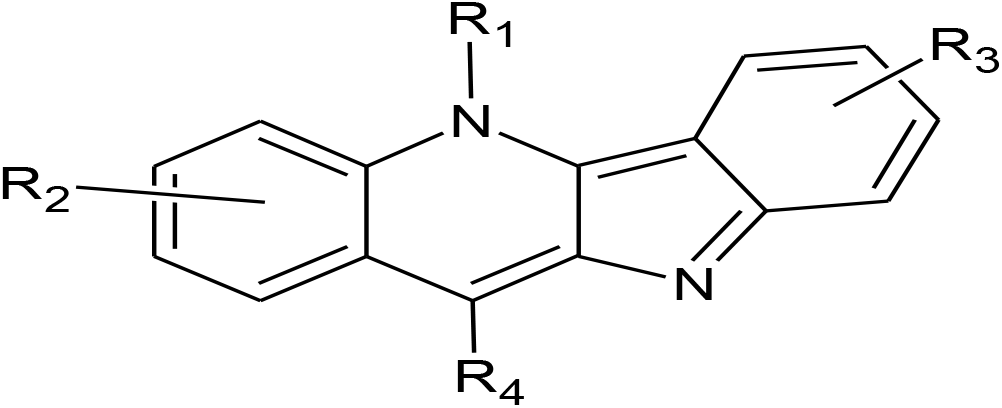
Structure of cryptolepine showing different functional groups at different positions as R_1_, R_2_, R_3_, R_4_

**Figure 4. 4:**
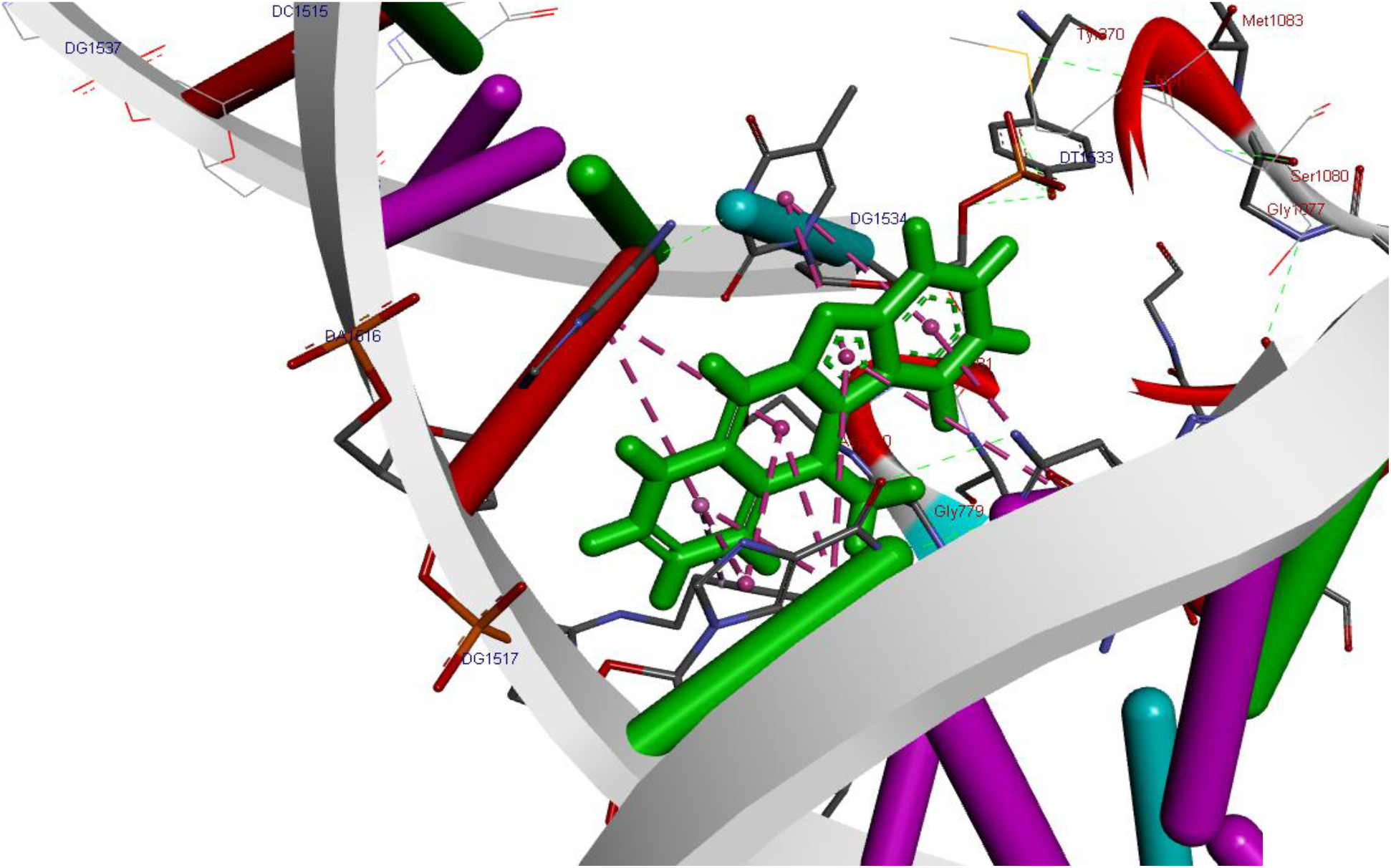
3D Structure showing the interaction of residues with cryptolepine in the active site of TOP2A

**Figure 4. 5:**
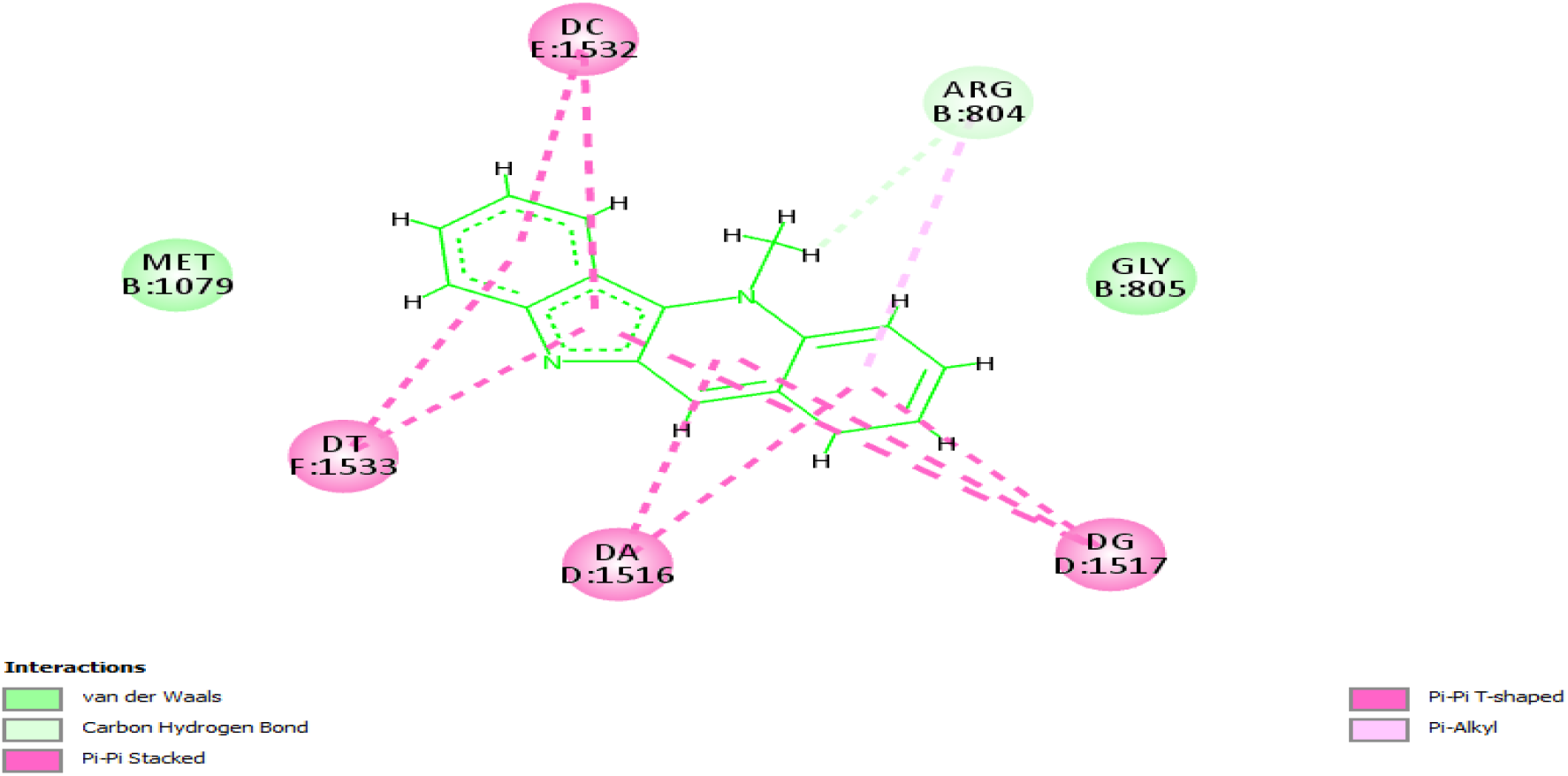
2D structure of cryptolepine showing the interacting residues with TOP2A

Cryptolepine interacts with the receptor TOP2A with residues such as Met 1079, Arg 804, Gly 805 and nucleotides dT 1533, dC 1532, dA 1516 and dG 1517. The residues Arg 804, Gly 805 and Met 1079 forms carbon hydrogen bond and van der waals interaction with the ligand, while the nucleotides dT 1533, dC 1532, dA 1516 and dC 1517 form pi-pi stacked, pi-pi T- shaped and pi- alkyl interactions. All these interactions result in a total binding energy of cryptolepine with receptor as −6.09 kcal/mol.

### 4.4 Virtual screening of cryptolepine Library

Virtual screening of the combinatorial library of 31114 molecules was done to identify potent inhibitor of TOP2A. Based on scoring function and strong interactions with receptor molecules five best ligands were selected. The top five molecules and their respective binding energies in kcal/mol are represented by ligand number 8618, 907, 145, 16755, 8186 as shown in the table 4.1. All the top 5 compounds along with the standard are superimposed in the binding pocket of TOP2A as shown in Figure 4.6.

**Figure 4. 6:**
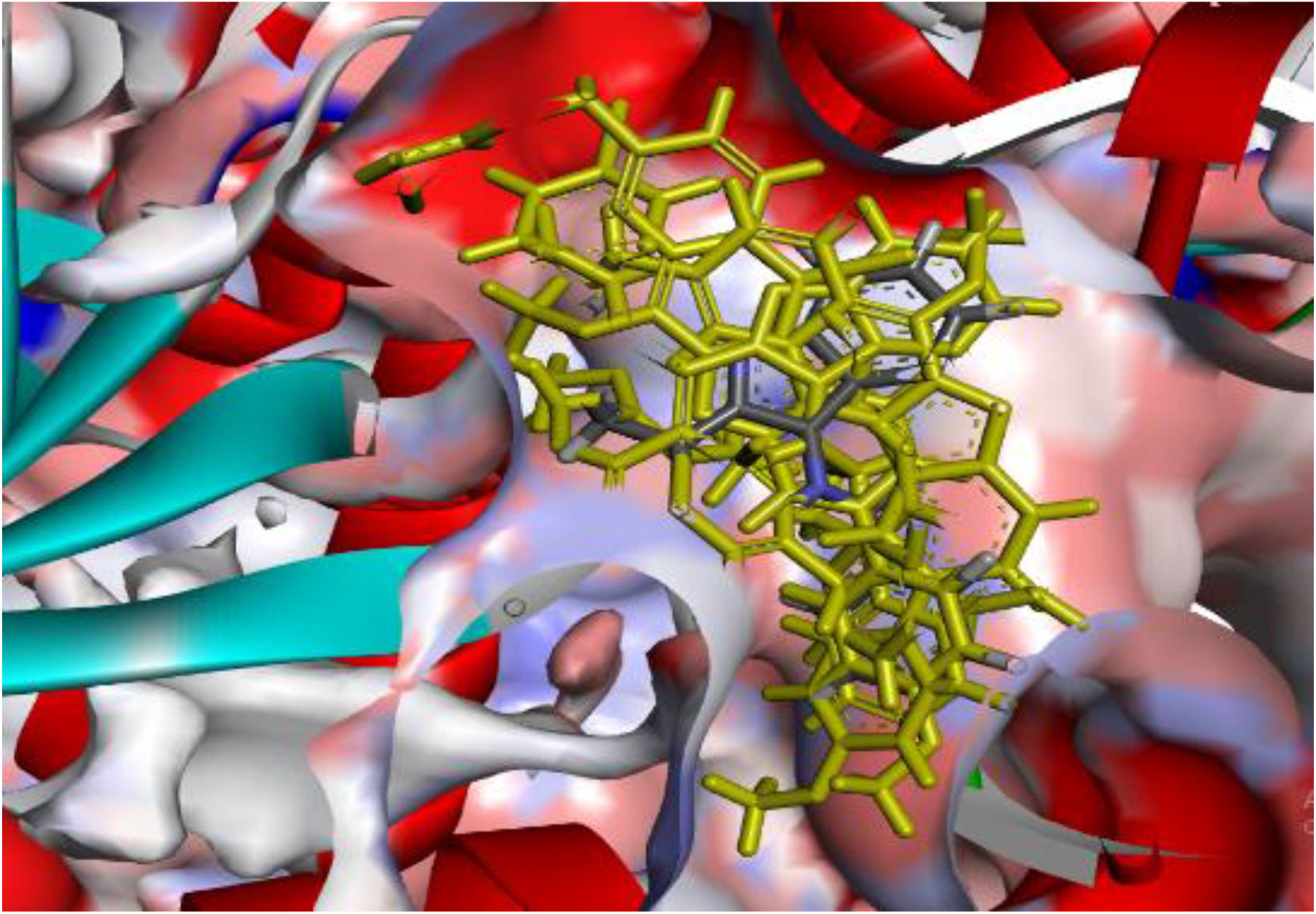
Docked poses of the top five selected derivatives and the standard cryptolepine all superimposed. The standard is depicted by cpk color and the five derivatives are shown as yellow colored sticks.

**Table 4.1:**
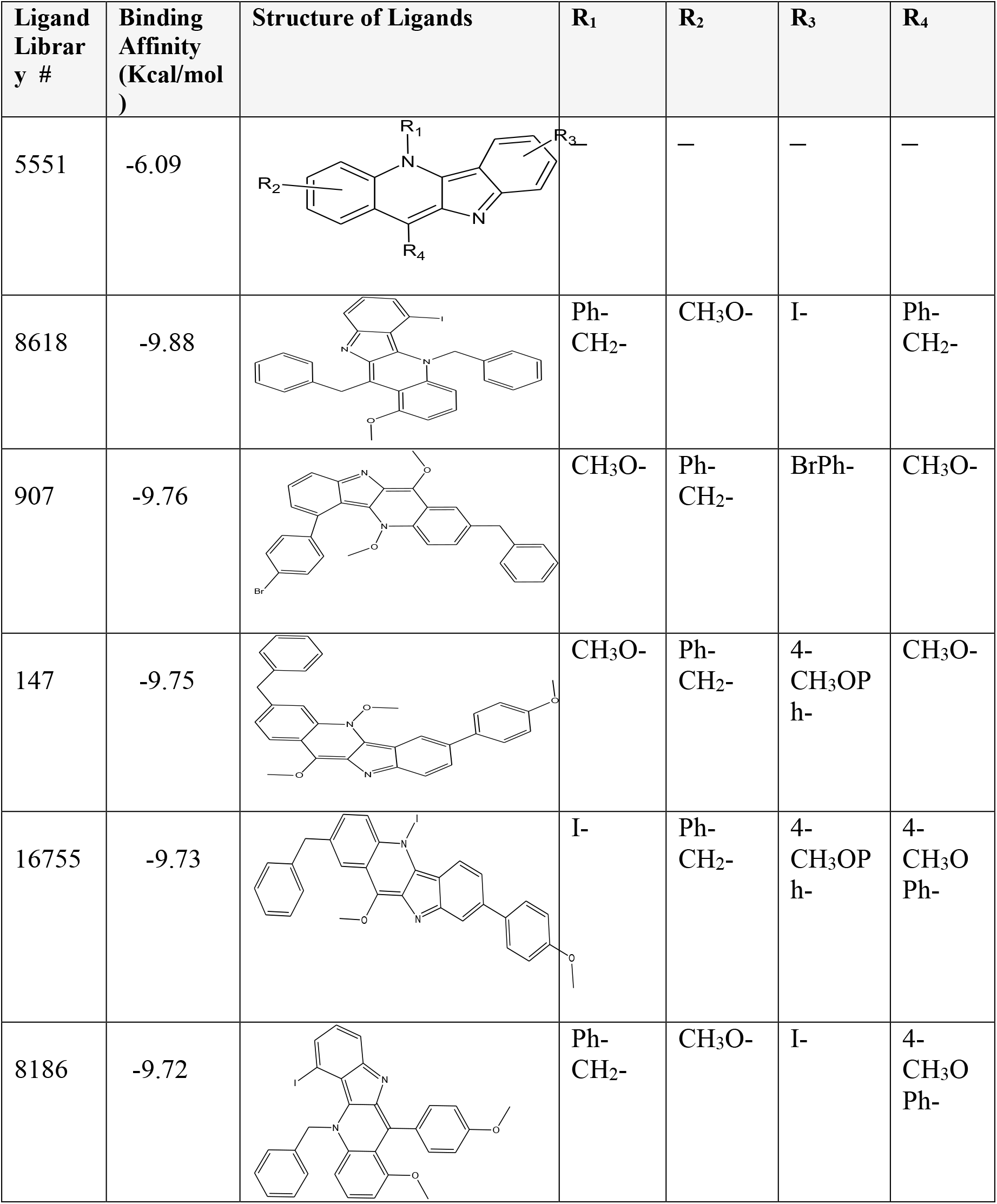
MOE docking score of top 5 derivatives for TOP2A

### 4.5 Detailed Interaction Analysis of the Docked Ligands in the Binding Pocket of Receptor TOP2A

Top 5 docked derivatives of cryptolepine were selected based on their scoring function to evaluate the binding interactions and energies for the receptor TOP2A. The details of various interactions and energy values of all the selected derivatives are given below.

#### 4.5 .1 Derivative no. 8618

Ligand no. 8618 interacts with TOP2A by amino acid residues Met 1079, Gly 1077, Ser 781, Tyr 370, Gly 779, Leu 803, Gly 805, Asp 780 and Arg 804 while nucleotides involved are dG 1517, dT 1533, dC 1532 dG 1534 and dA 1516. Ph_CH2 (Benzyl group) attached at position R_1_ form pi-pi stacked interaction with dG 1517, OCH_3_ group attached with at position R_2_ form carbon hydrogen bond with Asp 780, van der waals interactions with Gly 805 and pi-alkyl interaction with Arg 804. Iodide (I) attached at position R_3_ forms pi-alkyl and pi-anion interactions with dT 1533. Benzyl group attached at position R_4_ forms van der waals interactions with amino acid residues such as Ser 781, Tyr 370, Gly 779, Leu 803 and with the nucleotide dG 1534. Due to these extra interactions formed by ligand no. 8618 with TOP2A the binding energy is lowered from −6.09 kcal/mol to −9.88kcal/mol which makes it more stable than cryptolepine with the TOP2A receptor.

**Figure 4. 7:**
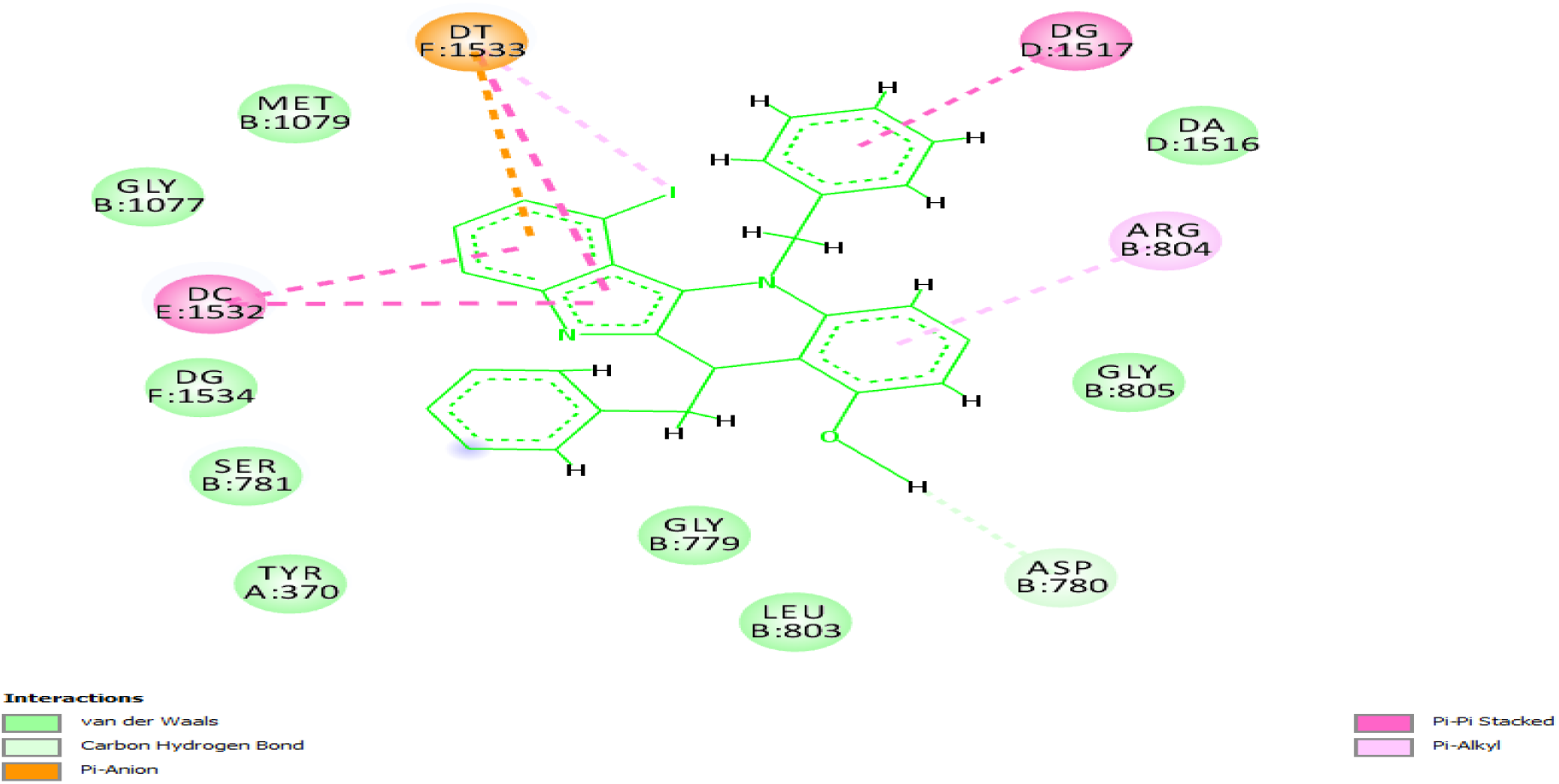
2D structure of ligand 8618 binding pocket green dotted line shows hydrogen bond while other lines show hydrophobic interactions

**Figure 4.8:**
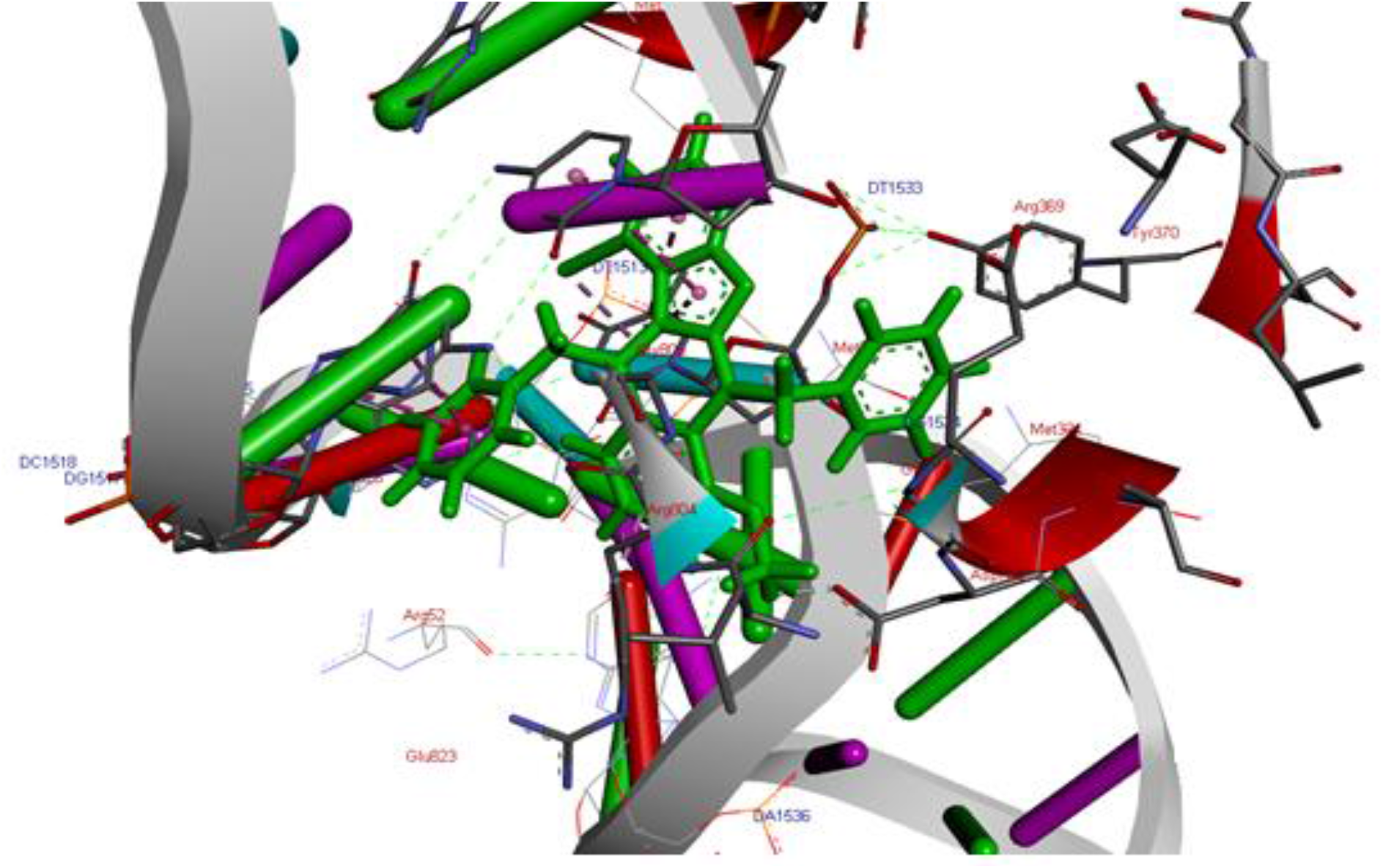
Interacting residues within the binding pocket of ligand no.8618 shows contacts with TOP2A.

#### 4.6.2 Derivative no. 907

Ligand no. 907 interacts with TOP2A by residues such as Ser 781, Arg 804, Asp 780, Try 370, Gly 805, Gly 779, Leu 803, Glu 778 and Asp 858 while nucleotides involved include dA 1516, dG 1534, dG 1517, dT 1533 and dC 1532. OCH_3_ at positon R_1_ forms van der waals interaction with Gly 779 and pi donar hydrogen bond with the nucleotide dC 1532. OCH_3_ (methoxy) group attached at position R_4_ forms carbon hydrogen bond with Asp 780. 4-Br Ph (bromophenol) group attached at position R_3_ forms van der waals interactions with amino acid Glu 778 and carbon hydrogen bond with nucleotide dC 1532. Benzyl group attached at position R_2_ forms pi-pi stacked, pi-pi T-shaped, amide-pi-stacked with nucleotides dT 1533, dG 1517 and van der waals interaction with dA 1516. Ligand no 907 can be considered better inhibitor than cryptolepine because the number of non-bonded interactions of the ligand no. 907 are more than cryptolepine taken as parent compound that’s why it has better binding affinity for the receptor than cryptolepine itself. All the hydrophobic interactions are responsible for lower binding energy. The binding energy for this is lowered to −9.76 kcal/mol from −6.09 kcal/mol (cryptolepine).

**Figure 4. 9:**
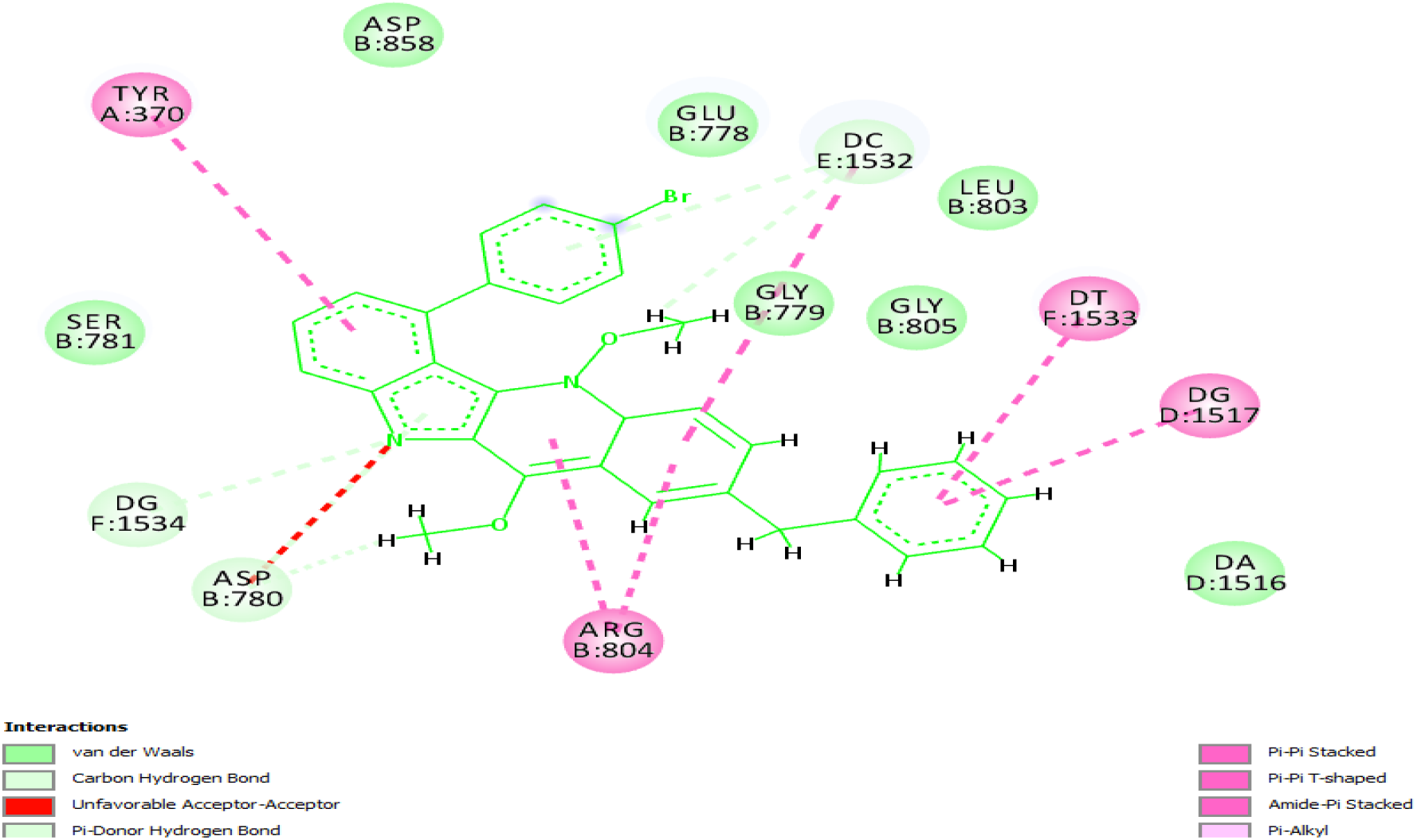
2D structure of ligand no. 907 represents dotted lines for hydrophobic contacts.

**Figure 4. 10:**
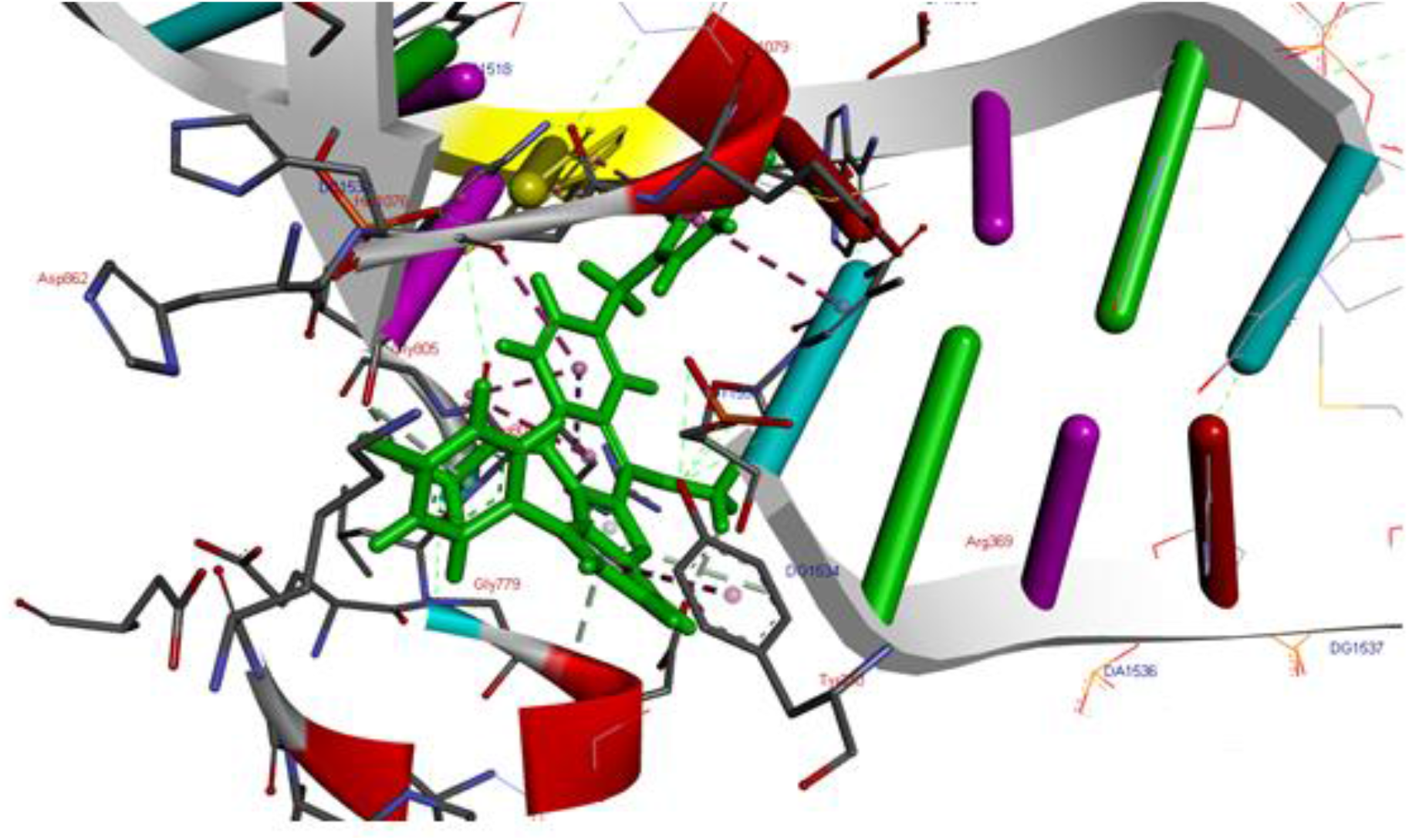
3D structure shows interacting residues within the binding pocket of ligand no 907

#### 4.7.3 Derivative no. 147

Ligand no 147 interacts with the TOP2A by amino acid residues such as Gly 1077, Met 1079, Ser 1080, Met 1083, Glu 823, Gly 805, Gly 779, Asp 780, Ser 781, Arg 804, Tyr 370 and with the double-stranded DNA such as dC 1532, dT 1533, dA 1516, dG 1531, dG 1534 and dG 1517. OCH_3_ group attached at position R_1_ forms van der waals interactions with Gly 779. Benzyl group attached at position R_2_ forms van der waals interactions with amino acid residues Asp 780, Ser 781, Tyr 370 and with the nucleotide dG 1534. 4-CH_3_OPh attached at position R_3_ forms pi-alkyl with amino acid Arg 804 and pi-pi stacked with the nucleotide dG 1517. OCH_3_ attached at position R_4_ forms pi-donor hydrogen bond with dC 1532 and van der waals interactions with nucleotide dG 1531 and with the amino acid residues Ser 1080, Met 1079. These extra interactions formed by ligand 147 make it more stable than cryptolepine to have more strong and stable interaction with the receptor. The binding energy for ligand no. 147 is lowered to −9.75 kcal/mol due to attachment of functional groups.

**Figure 4. 11:**
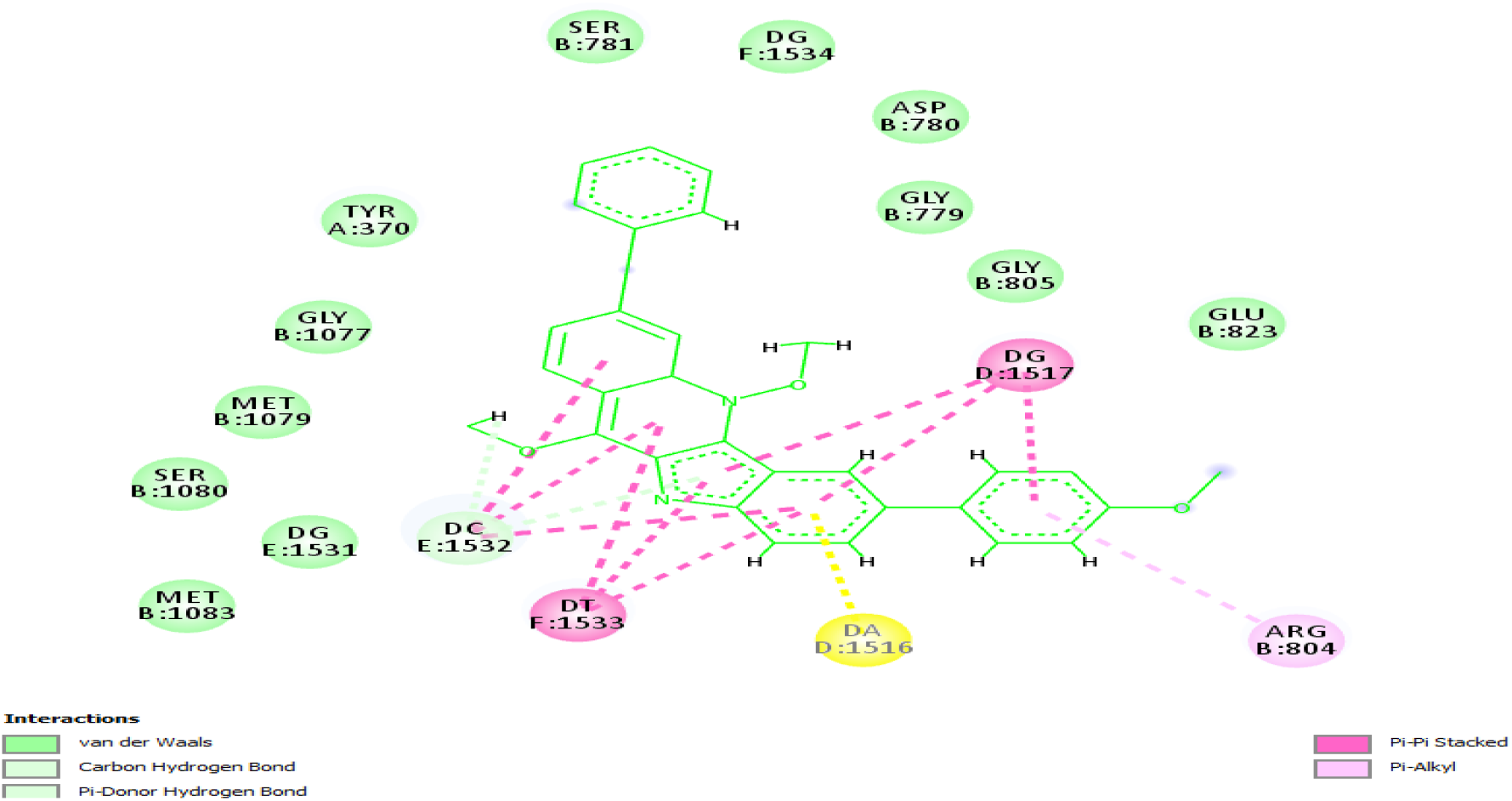
2D structure of ligand 147 showing the interactions by dotted lines.

**Figure 4. 12:**
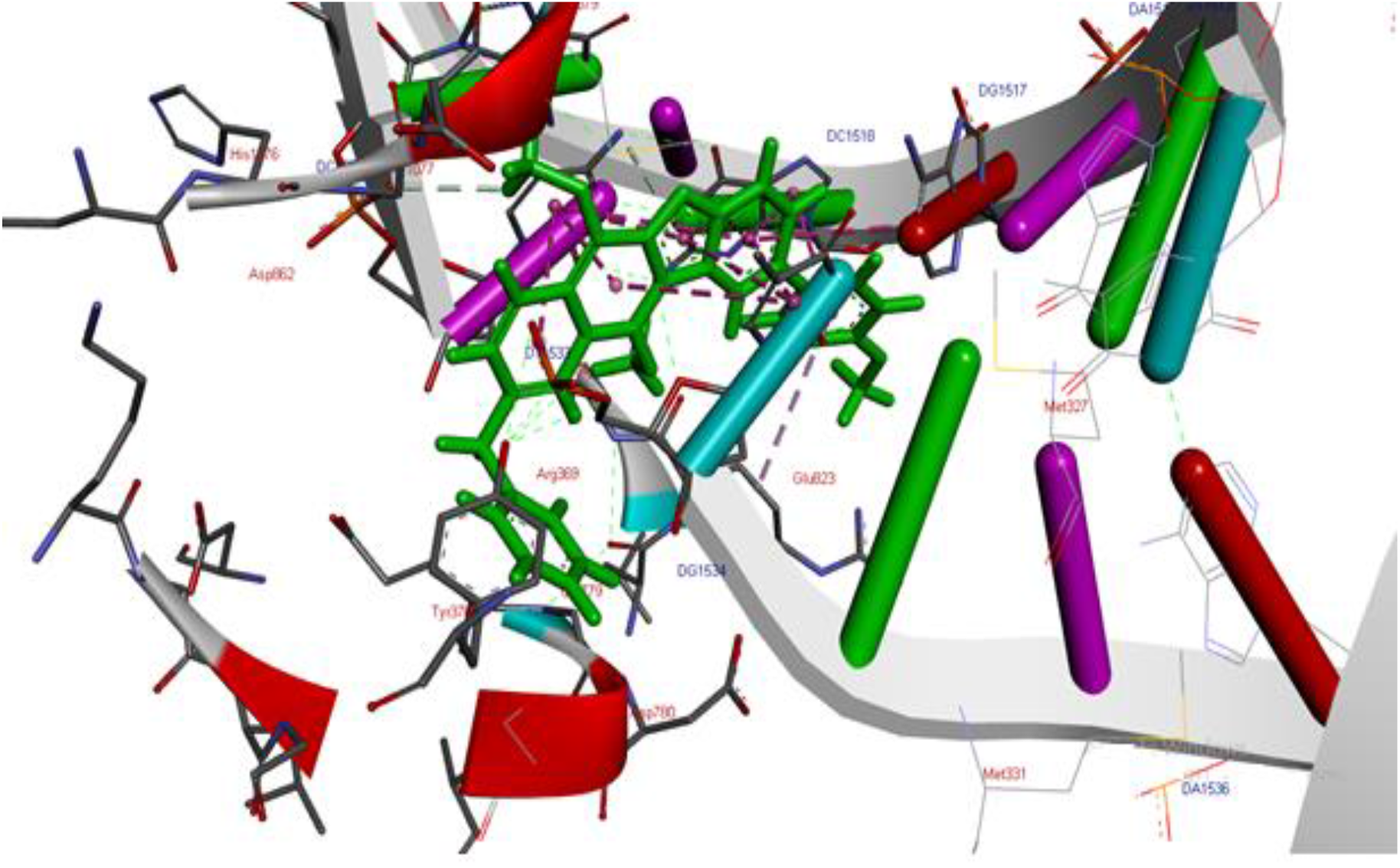
3D structure shows the interacting residues within the binding pocket of ligand no. 147 within TOP2A

#### 4.8.4 Derivative no. 16755

Ligand no 16755 interacts with the residues such as Met 1079, Gly 1077, His 1076, Lys 931, Gly 932, Ser 781, Leu 933, Ala 782, Gly 779, Asp 858, Gly 805, Arg 804, Gly 934, Glu 778, and Tyr 370 while nucleotides include are dC 1532, dT 1533, dG 1517 and dA 1516. Benzyl group attached at position R_2_ forms carbon hydrogen bond with Gly 934 and van der waals interactions with amino acid residues Ser 781, Gly 932, Tyr370 and Gly 931. CH_3_OPh attached at position R_3_ forms pi-donor hydrogen bond and pi-pi stacked interactions with dG 1517 and van der waals interactions with dA 1516. OCH_3_ group attached at position R_4_ forms carbon hydrogen bond with Glu 778 and van der waals interactions with Gly 779 and Asp 858. All the interactions lower the binding energy from −6.09 kcal/mol to −9.73 kcal/mol which makes it best fit within the binding pocket.

**Figure 4. 13:**
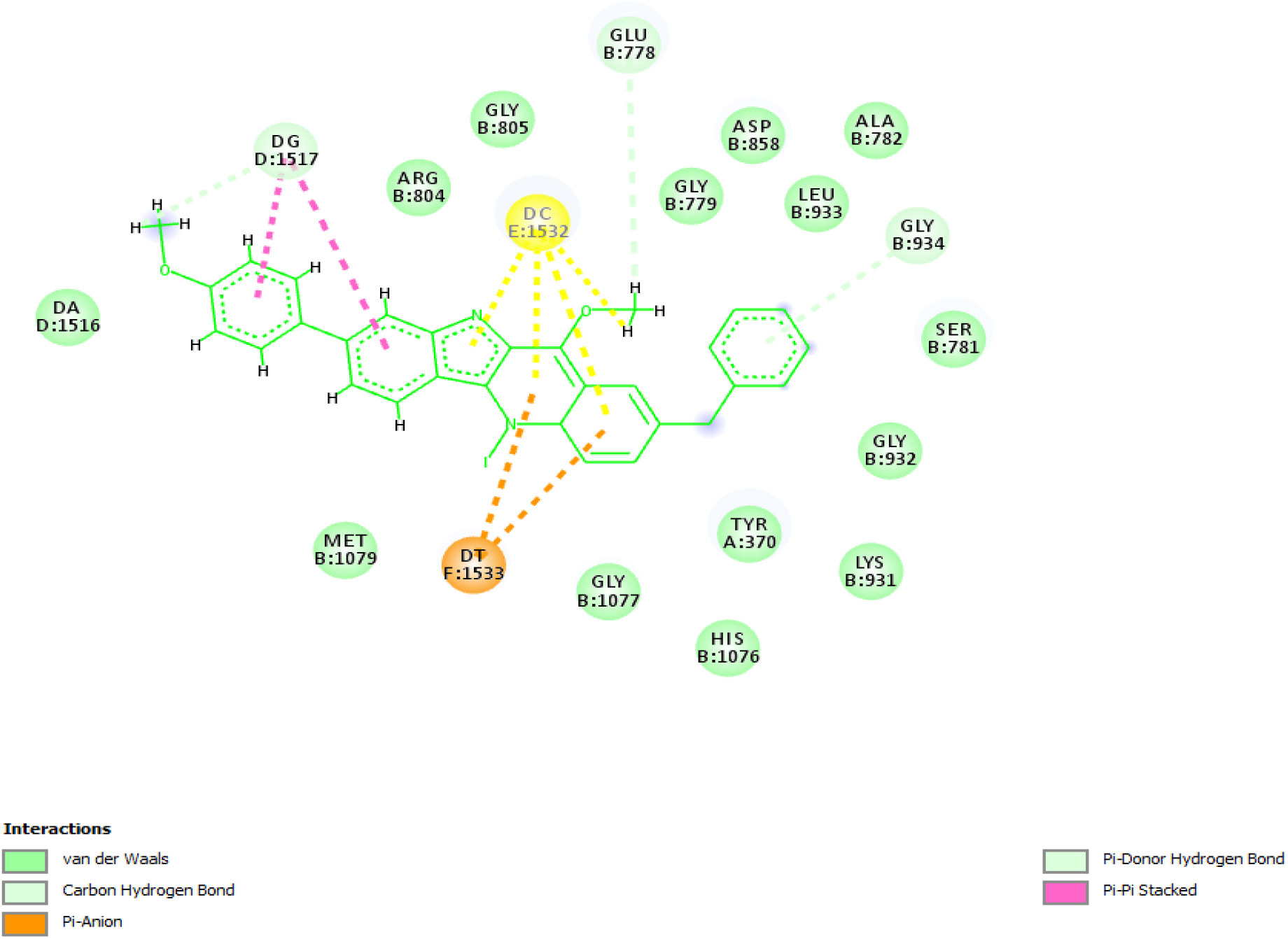
2D structure of ligand no 16755 shows hydrophobic contacts by dotted lines.

**Figure 4. 14:**
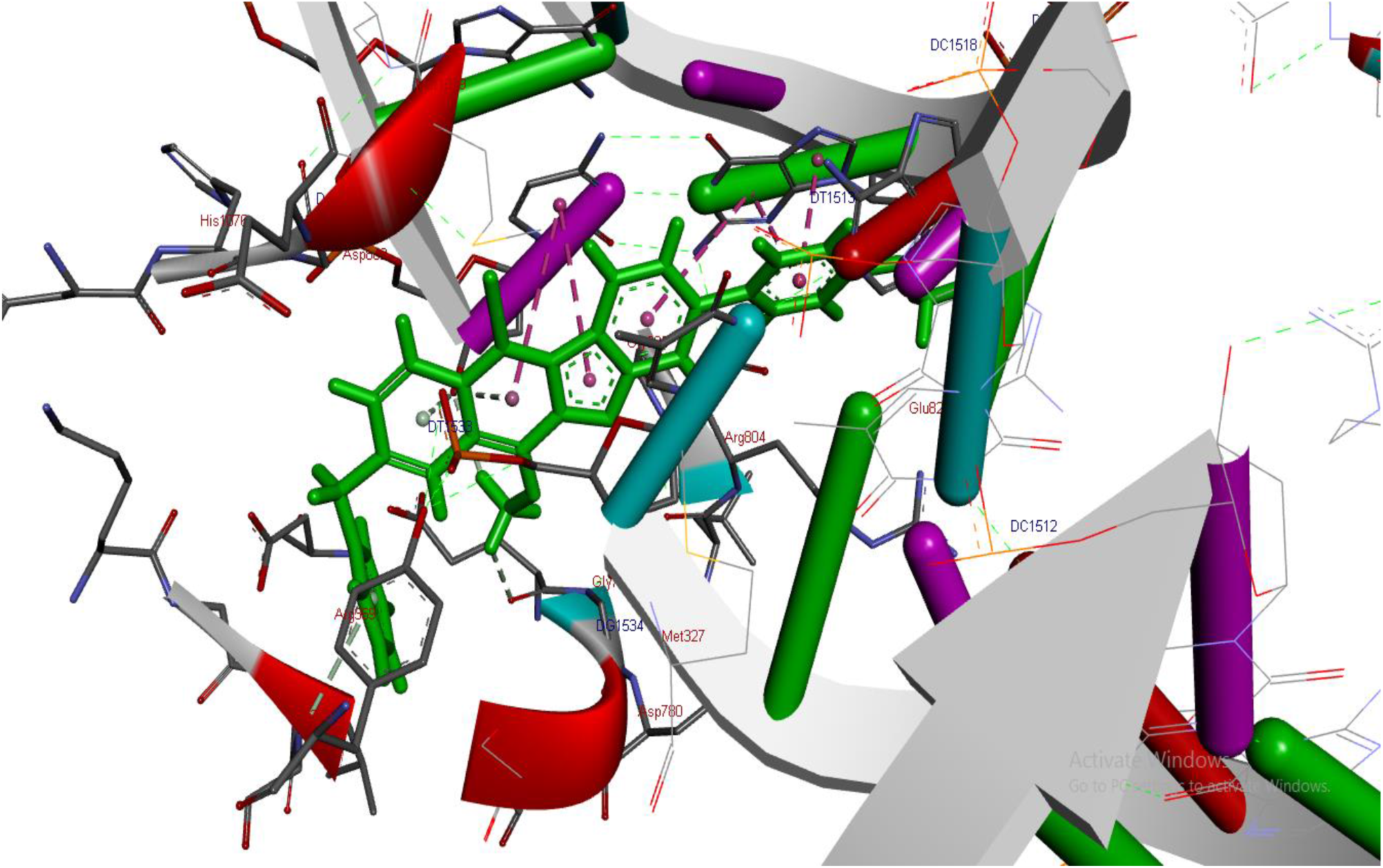
3D structure representing interacting residues within the binding pocket of ligand no 16755.

#### 4.9.5 Derivative no. 8186

All the interactions are responsible for best binding affinity and lower binding energy which is −9.72 kcal/mol for ligand no. 8186. The type of residues of TOP2A with which ligand 8186 interact are Met 1083, Ser 1080, Gly 1077, Leu 933, Gly 932, Ala 782, Asp 780, Gly 805, Arg 804, Gly 779, Leu 803, Met 1079, Ser 781 and Glu 778 while nucleotides interact by dA 1516, dG 1517, dT 1533, dG 1534 and dC 1532. Benzyl group attached at position R_1_ forms pi-pi stacked and pi-pi T-shaped interactions with dA 1516 and dG 1517 and pi-alkyl interactions with Arg 804. Methoxy group attached at position R_2_ forms carbon hydrogen bond with amino acid residues Leu 803 and Gly 779 and van der waals interactions with Gly 805. I group attached at position R_3_ forms pi-alkyl with dT 1533. CH_3_OPh group attached at position R_4_ forms carbon hydrogen interactions with Ser 781 and Gu 778 and van der waals interactions with Gly 932, and Ala 782. All the interactions are responsible for best binding affinity and lower binding energy which is −9.72 kcal/mol for ligand no. 8186.

**Figure 4. 15:**
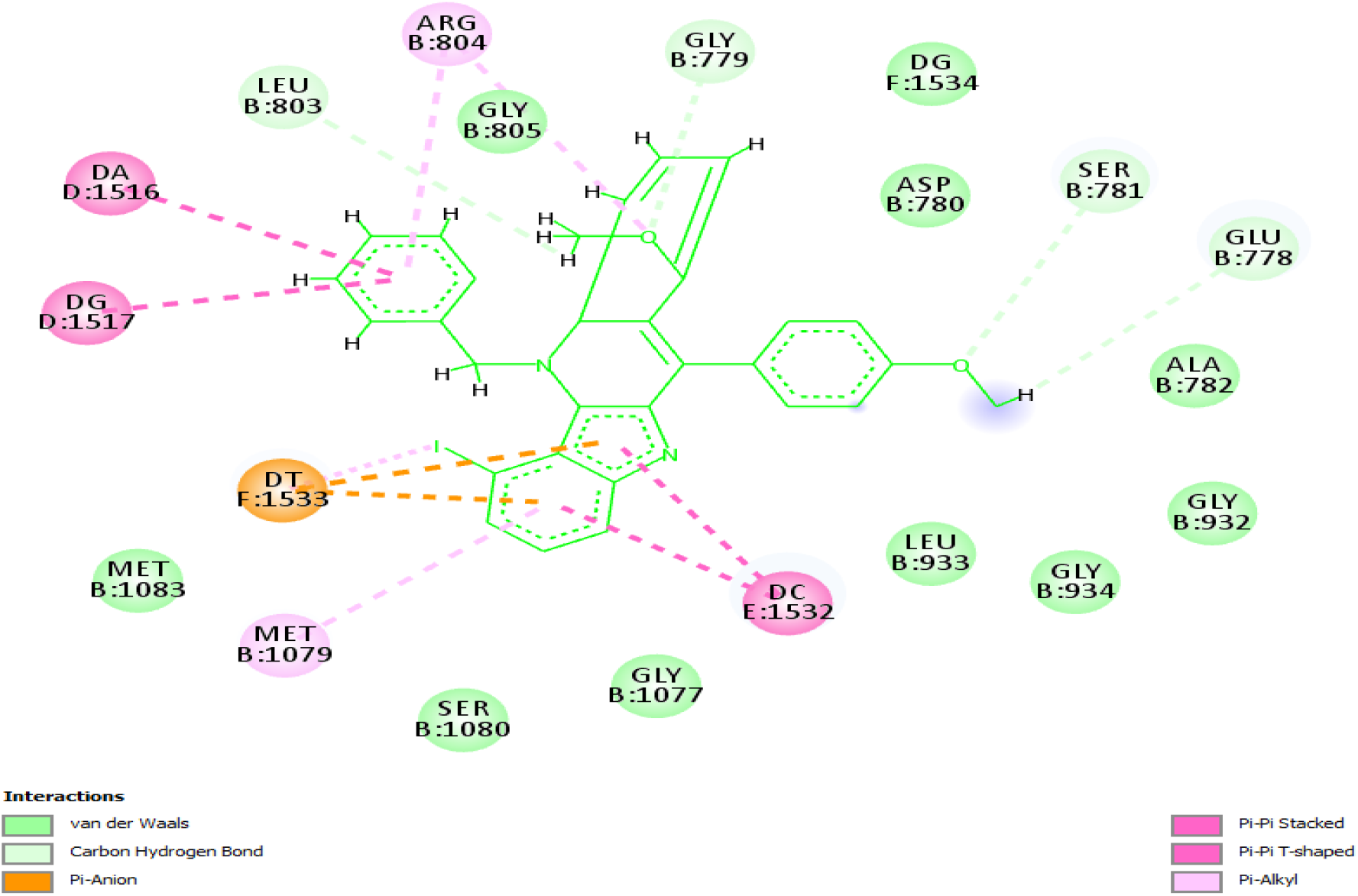
2D structure of ligand no. 8186 representing different hydrophobic interactions

**Figure 4. 16:**
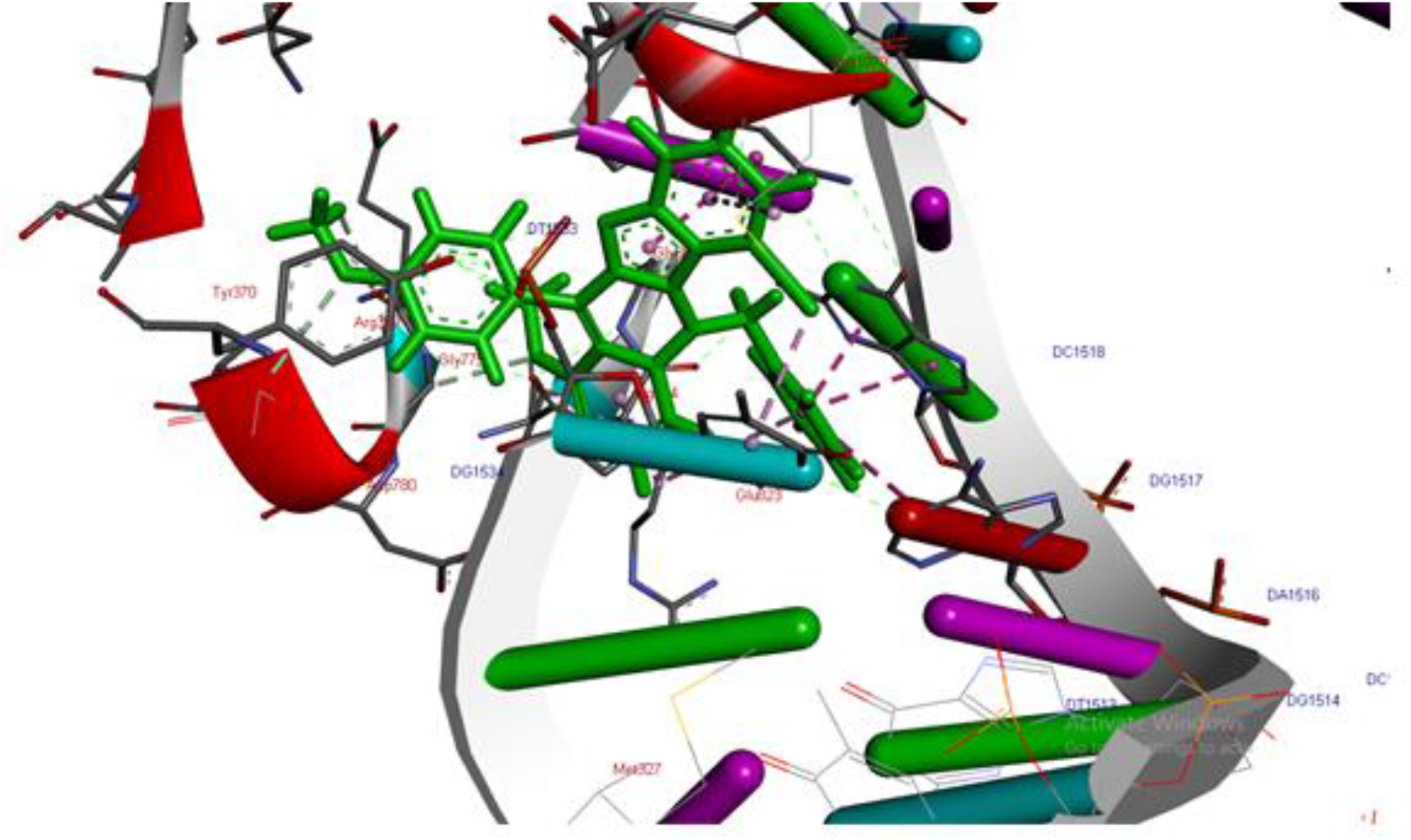
3D structure showing interacting residues within the binding pocket of ligand no. 8186

## Conclusion

TOP2A is a key enzyme in human dividing cells and its activity is greatly enhanced in dividing cells, therefore, it can be considered as a good target for designing new anticancer drugs. Computer-aided drug designing and docking analysis plays a key role in designing new inhibitors for a particular protein. The inhibitory effect of the drug toward the target protein can be determined by calculating their binding energies and can be explained by observing the ligand interaction with various amino acids residues of the binding pocket of the target protein.

The present work includes the virtual screening of 31114 derivatives of cryptolepine by molecular docking study. Top five inhibitors of TOP2A were selected based on good binding energy. The study reveals that ligand no. 8618, 907, 147, 16755 and 8186 are the more potent ligands among the 31114 ligands since they form number of hydrophobic interactions with the target receptor thereby lowering the predicted binding energy. The binding energies of the Ligand no. 8618, 907, 147, 16755 and 8186 are predicted as −9.88kcal/mol, −9.76 kcal/mol, −9.73 kcal/mol, −9.72 kcal/mol and −9.72 kcal/mol respectively. All the known inhibitors of TOP2A were also docked into the binding pocket and it was observed that the selected ligands from the cryptolepine derivatives library has better binding energies and more favorable interactions with receptor.

So, it is concluded that cryptolepine can be modified to a number of best inhibitors to enhance the potency and lessen the side effects of existing drugs targeting human TOP2A.

We propose a set of derivatives of the cryptolepine that are predicted to have better binding affinity towards TOP2A. All these selected ligands fit very well into the binding pocket with minimum solvent accessible surface area (SASA). The substituents putted on various positions of cryptolepine in the selected ligands were chosen for the fact that they can be easily incorporated into the starting compounds required for the total synthesis of cryptolepine. The total synthesis of cryptolepine has been reported with different synthetic schemes [17].

